# Structure-preserving fixation allows Scanning Electron Microscopy to reveal biofilm microstructure and interactions with immune cells

**DOI:** 10.1101/2023.10.25.564022

**Authors:** Marilyn Wells, Michelle Mikesh, Vernita Gordon

## Abstract

*Pseudomonas aeruginosa* is a pathogen that forms robust biofilms which are commonly associated with chronic infections and cannot be successfully cleared by the immune system. Neutrophils, the most common white blood cells, target infections with pathogen-killing mechanisms that are rendered largely ineffective by the protective physicochemical structure of a biofilm. Visualization of the complex interactions between immune cells and biofilms will advance understanding of how biofilms evade the immune system and could aid in developing treatment methods that promote immune clearance with minimal harm to the host. Scanning electron microscopy (SEM) distinguishes itself as a powerful, high-resolution tool for obtaining strikingly clear and detailed topographical images. However, taking full advantage of SEM’s potential for high-resolution imaging requires that the fixation process simultaneously preserve both intricate biofilm architecture and the morphologies and structural signatures characterizing neutrophils responses at an infection site. Standard aldehyde-based fixation techniques result in significant loss of biofilm matrix material and morphologies of responding immune cells, thereby obscuring the details of immune interactions with the biofilm matrix. Here we show an improved fixation technique using the cationic dye alcian blue to preserve and visualize neutrophil interactions with the three-dimensional architecture of *P. aeruginosa* biofilms.

## Introduction

Biofilms are communities of microbes encased in a self-produced matrix of extracellular polymeric substances (EPS). Compared with their planktonic counterparts, biofilm microbes have enhanced resistance to antibiotic treatment and immune clearance. The EPS matrix consists of polymers, proteins, nucleic acids, lipids, and additional material from the host environment [1-4]. Biofilms are the cause of up to 80% of persistent infections, notably in the lungs of patients with Cystic Fibrosis (CF), in chronic wounds, and on foreign surfaces in the body such as catheters or implants [5, 6]. All these cases frequently involve *Pseudomonas aeruginosa*, a common opportunistic human pathogen which forms robust biofilms and serves as a model organism for the study of biofilm infections [7, 8].

Current investigations aim to understand the interactions between biofilms and the immune system to develop treatment plans which result in successful immune clearance. Neutrophils, which are up to 70% of all white blood cells, serve as the first line of immune defense and are chemotactically recruited to an infection site [9]. Neutrophils employ three primary killing mechanisms: phagocytosis, degranulation, and Neutrophil Extracellular Traps (NETs). Phagocytosis involves the engulfment and internalization of microbes into a digestive vesicle called a phagolysosome, within which bacteria are subsequently killed and degraded [10, 11]. Degranulation is the localized release of preformed vesicles containing antimicrobial enzymes [12]. NETosis is the expulsion of webs of DNA to trap and kill nearby bacteria [13].

When faced with a biofilm infection that evades immune clearance, neutrophils’ primarily killing mechanisms are often rendered ineffective and may ultimately harm the host via poorly controlled release of factors intended to kill pathogens [9, 14]. For example, attempted phagocytosis of a target much larger than the size of a neutrophil results in ‘frustrated’ phagocytosis. Neutrophils spread themselves across the target but are unable to close and form a phagosome, causing a dangerous inflammatory response [15]. For another example, DNA released through NETosis can be taken up by the biofilm as a structural component of the matrix, further inhibiting successful immune clearance [4, 16]. Some bacterial species have even been shown to modulate neutrophil degranulation in their favor: inhibiting degranulation reduces neutrophils’ microbicidal effects, while inducing more can be harmful to the surrounding host environment [12].

Biofilms are heterogeneous structures whose composition and mechanical properties result in complex interactions with immune cells [12, 17-19]. Detailed visualization of these interactions is critical to the development of effective treatments to compromise biofilms and promote clearance by the host immune system. Scanning electron microscopy (SEM) is a powerful tool for obtaining detailed insight to guide these studies. However, what SEM can reveal about immune cells responding to biofilms and interacting with the biofilm structure is limited by how much of the original biofilm material and structure can be preserved through the fixation process. Classical microbiology techniques for SEM aim to preserve microbial cells using aldehyde-based fixatives such as glutaraldehyde or paraformaldehyde, both of which efficiently penetrate cells, cross-link proteins, and halt cellular processes [20-22]. Osmium tetroxide is commonly used as an electron contrast stain as well as a secondary fixative to retain lipids in tissue samples and cell membranes [23]. Newer methods of fixing biofilms have used cationic dyes to preserve the polysaccharide components of the EPS matrix, thereby allowing SEM to reveal more intricate details of the matrix structure than was possible using prior approaches [24]. Without these details, the role of particular EPS components on biofilm structure may largely be lost.

Here we demonstrate the use of a substantially enhanced fixation technique that allows SEM to visualize in high detail the interactions between neutrophils and *P. aeruginosa* biofilms. This includes the role of particular EPS components in biofilm microstructure, and the microstructure’s interaction with neutrophils, that is not preserved through previously-published fixation techniques. Our work here paves the way for future studies to gain critical insight into immune system’s pathogen-targeting mechanisms amidst the challenges created by biofilm formation.

## Methods

### Isolation of neutrophils

Work with neutrophils was approved by the Institutional Review Board at the University of Texas at Austin (Austin, TX) as Protocol No. 2021-00170. Neutrophils from healthy adult volunteer blood donors were isolated following a previously published protocol [25]. Blood was collected by venous puncture and drawn into lithium-heparin coated tubes (BD Vacutainer). Whole blood was mixed with an equal volume of a filter-sterilized (VWR) solution containing 3% dextran and 0.9% sodium (Sigma-Aldrich). Red blood cells were allowed to sediment and fall to the bottom of the tube. The resulting supernatant was centrifuged (Eppendorf 5810R, A-4-62 Rotor) for 10 minutes at 500 x g to separate the cells from plasma. The resulting pellet was resuspended in Hanks Buffered Salt Solution (HBSS) without calcium or magnesium (Gibco Laboratories), gently layered onto a Ficoll-Paque density gradient (Sigma), and centrifuged for 40 minutes at 400 x g. The supernatant was discarded and the remaining pellet suspended in deionized water to lyse any remaining red blood cells. The solution was stabilized by the addition of a filter-sterilized 1.8% NaCl solution. Following 5 minutes centrifugation at 500 x g, a pellet of neutrophils was resuspended in 800 μL HBSS with calcium and magnesium and 200 μL human serum (Sigma). Each 10 mL lithium-heparin coated tube of blood drawn yielded 1 mL of neutrophil-only solution. In experiments where more than one tube of blood was drawn, the resulting neutrophil solutions were thoroughly mixed by pipette.

### Growth media preparation

Liquid growth media was prepared by adding 30 g LB broth powder (Fisher) to 750 mL deionized water. Agar media was prepared by combining 20g of LB agar powder (Fisher) to 500 mL deionized water. Both forms of media were sterilized with a liquid autoclave cycle. The agar media was allowed to cool, stirring, for 30 minutes and immediately poured into sterile petri dishes (Fisher). Agar plates were allowed to solidify overnight at room temperature.

### Biofilm growth conditions

The bacterial strains used in this study are PA01 WT and PA01 mutant PW2387 (genotype mucA-A05:ISphoA/hah) [26], which overproduces the polymer alginate. Frozen bacterial stock was streaked onto a sterile LB agar plate and allowed to grow overnight at 37 °C. A single colony was selected and cultured in 4mL of fresh LB broth and grown shaking overnight at 37 °C. Biofilms were grown on squares of ACLAR®(Ted Pella) by submerging the substrate in 1mL of overnight culture and allowing them to grow statically overnight. Excess culture was removed and samples transferred to petri dishes for SEM fixation. For studies with neutrophils, excess culture was removed and 200 μL of neutrophil solution was added and allowed to incubate for 1 hour at 37 °C. Excess solution was removed and samples transferred to petri dishes for fixation.

### Fixation for SEM

Fixatives and post-fixatives for the three procedures used are listed in Table 1. As a standard fix, we use 2% glutaraldehyde followed by 1% osmium tetroxide. An improved standard fix includes the addition of 2% paraformaldehyde to the standard initial fixative. We use an enhanced fix, adapted from methods described in [24], which includes 2% glutaraldehyde, 2% paraformaldehyde, and 0.15% alcian blue in the initial fixative, followed by 1% osmium tetroxide and 1% tannic acid in the post-fixative.

**Table 1.** Details of initial and post-fixatives used in each of the three tested fixation techniques.

|  | **Standard Fix** | **Improved Standard Fix** | **Enhanced Fix (adapted from [24])** |
| --- | --- | --- | --- |
| *Initial fixative (in 0.1M Cacodylate buffer at pH 7-7.4)* | 2% glutaraldehyde | 2% glutaraldehyde + 2% paraformaldehyde | 2% glutaraldehyde + 2% paraformaldehyde + 0.15% alcian blue |
| *Post-fixative* | 1% osmium tetroxide | 1% osmium tetroxide | 1% osmium tetroxide + 1% tannic acid |

Samples were treated at room temperature with an initial fixative, covered, and allowed to fix overnight for approximately 16 hours (Glutaraldehyde and Paraformaldehyde, Electron Microscopy Sciences; Alcian Blue, Sigma). The samples were washed thrice with 0.1M sodium cacodylate (Electron Microscopy Sciences), with 10 minutes per wash. The buffer was removed and samples stained with post-fixative for 3 hours (Osmium tetroxide, Ted Pella; Tannic Acid, Electron Microscopy Sciences). After washing with distilled water, biofilms were dehydrated with graded alcohols for 15 minutes per step, 5 minutes in 1:1 absolute ethanol (Thermo) to hexamethyldisilazane (HMDS, Ted Pella), and 5 minutes in 100% HMDS. Samples were air-dried for at least 30 minutes. Specimens were mounted to clean SEM stubs using double-sided conductive tape (Ted Pella), and then coated with 12 nm platinum/palladium using a Cressington 208HR sputter coater (Ted Pella).

### Imaging

Samples were imaged under high vacuum with a Zeiss Supra field emission SEM using an SE2 detector and a 5kV accelerating voltage. All reported experiments were performed in triplicate, and presented figures are representative of each experimental condition.

## Results and Discussion

### Glutaraldehyde and paraformaldehyde fixatives (Standard and Improved Standard fixes) do not preserve 3-D biofilm structure or EPS

As illustrative examples of biofilm types, we use biofilms grown from two lab strains of *P. aeruginosa*, PA01 wild type (denoted WT) and mutant PW2387 (genotype mucA-A05:ISphoA/hah) [26] (denoted Alginate+). Alginate+ overexpresses the anionic polysaccharide alginate, whose production in CF airways has been associated with infection persistence and poor clinical condition of patients [27, 28]. A standard glutaraldehyde-based fixative (“Standard Fix” in Table 1) is commonly used to preserve proteins [22]. Glutaraldehyde molecules each possess two aldehyde groups separated by a chain of 3 methylene bridges. Since cross-linking can occur through both aldehyde groups, glutaraldehyde has high bonding potential with proteins [29], but fails to preserve carbohydrates, leading to cell shrinkage during graded-alcohol dehydration. SEM imaging of biofilms fixed with glutaraldehyde shows the biofilms as two-dimensional structures composed of a single layer of cells, and many cells appear to have collapsed during preservation (Figure 1A,D).

**Figure 1.**
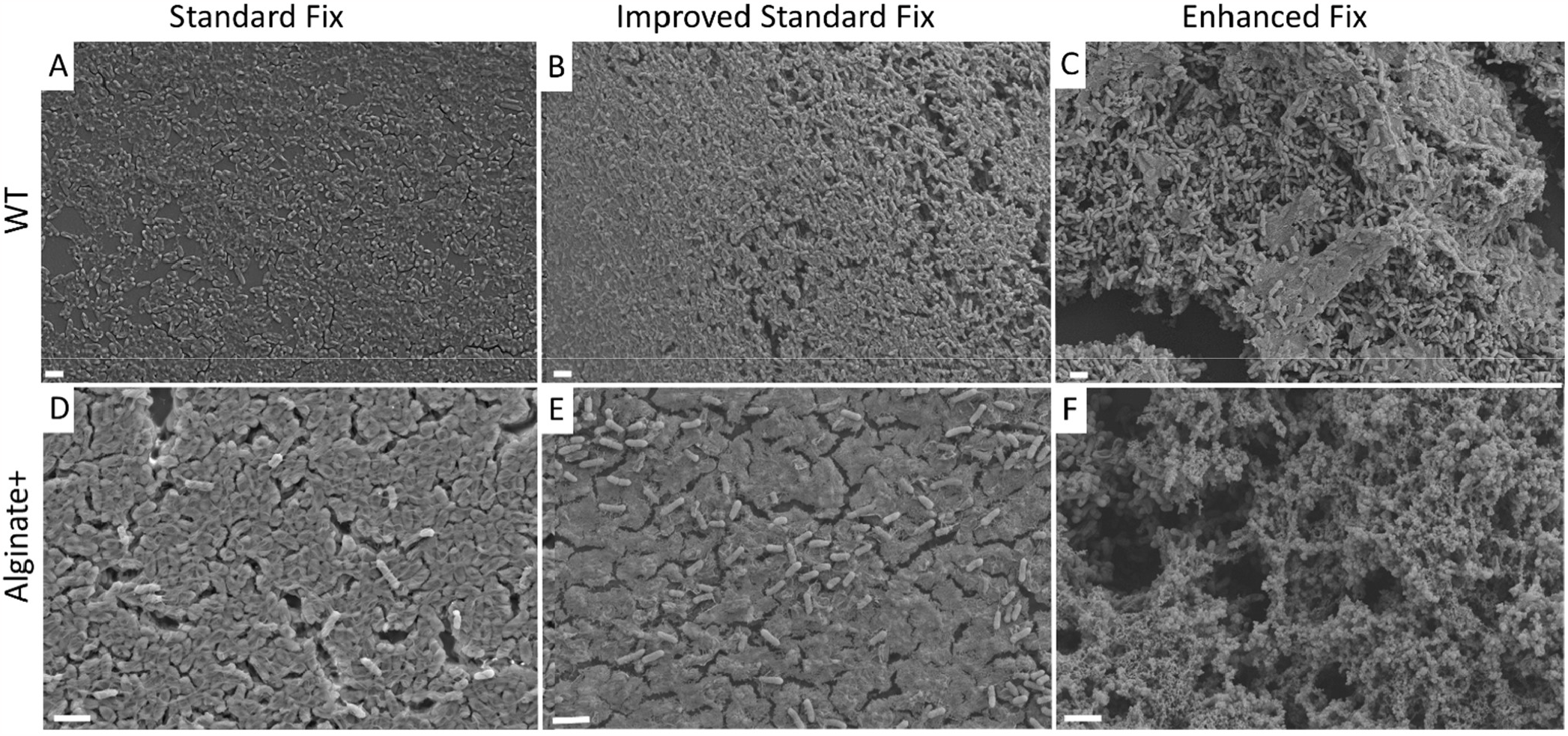
Representative SEM images of WT(A-C) and Alginate+ (D-F) *P. aeruginosa* biofilms preserved with 3 fixation methods. The Standard Fix used 2% glutaraldehyde as the initial fixative. The Improved Standard Fix used 2% glutaraldehyde and 2% paraformaldehyde as the initial fixative. Both the Standard and Improved Standard fixes used 1% osmium tetroxide as post-fixative. The Enhanced Fix used 2% glutaraldehyde, 2% paraformaldehyde, and 0.15% alcian blue as the initial fixative, with 1% osmium tetroxide and 1% tannic acid as post-fixative. Experiments were performed in biological triplicate. Scale bars represent 2µm.

The combination of glutaraldehyde and paraformaldehyde (“Improved Standard Fix” in Table 1) resulted in slightly higher retention of the WT biofilm’s EPS and its three-dimensional structure than did glutaraldehyde alone (Figure 1A,B), while Alginate+ biofilms did not appear to be significantly better preserved (Figure 1D,E). We speculate that this marginal difference may be connected to the fact that covalent cross-linking with cdrA is structurally important in Psl-dominated WT biofilms, which contain anionic, cationic, and neutral matrix polymers, whereas the matrices of Alginate+ biofilms are anionic and un-crosslinked.

### Enhanced fixation preserves polymeric substances and 3D structures

Cationic dyes have been used to improve EPS retention because they form ionic bonds with anionic polysaccharides present in the biofilm matrix [24, 30, 31]. To assess the effect of cationic dyes on preservation of *P. aeruginosa* matrix material, we add alcian blue to our initial fixative and tannic acid to the post-fixative (“Enhanced Fix” in Table 1). Tannic acid has been shown to act as a mordant between heavy metals and osmium-treated cellular structures [32, 33]. The increase in density of membrane components <u>improves contrast</u>, thereby <u>enhancing visualization</u> of intercellular connections and decreasing the amount of cell shrinkage during dehydration [24]. We found a dramatic improvement in EPS preservation with the addition of alcian blue and tannic acid (Figure 1C,F) when compared to the glutaraldehyde/paraformaldehyde combination or to glutaraldehyde alone. Increasing the amount of retained matrix material will enable work to more thoroughly visualize and analyze the three-dimensional architecture caused by EPS components.

Notably, SEM imaging of samples fixed using our improved technique qualitatively shows that mucoid biofilms are composed of a high volume fraction of alginate (Figure 1F), while this information was entirely lost to SEM imaging of samples prepared using the other two fixation methods. Hence, we see that the inclusion of cationic dyes in SEM sample preparation is critical for capturing an accurate picture of the biofilm matrix, particularly when the matrix is dominated by anionic polysaccharides.

Since *P. aeruginosa* infections in CF airways are largely dominated by alginate [7, 34-38], and since chronic inflammation caused by the immune response to biofilm infections is the major cause of lung damage for CF patients, we now turn our focus to the Alginate+ biofilms and their interactions with neutrophils.

### Enhanced fix allows detailed visualization of neutrophil killing mechanisms

Neutrophils exhibit a host of antimicrobial mechanisms and behaviors which may depend on the physicochemical properties of their target [9, 39]. To assess the effect of each fixation method on preservation of biofilm-neutrophil interactions, we allowed freshly-isolated human neutrophils to incubate with Alginate+ biofilms for 1 hour immediately prior to fixation and preparation for SEM imaging. Each of the three fixation methods listed in Table 1 were evaluated. While some biofilm material is preserved by each method, more complex structures and volumes were retained with the enhanced fix; this allowed observation of three-dimensional biofilm matrix structure and the manifestation of neutrophil activities in response to a biofilm infection that could not be seen using the other two methods (Figure 2).

**Figure 2.**
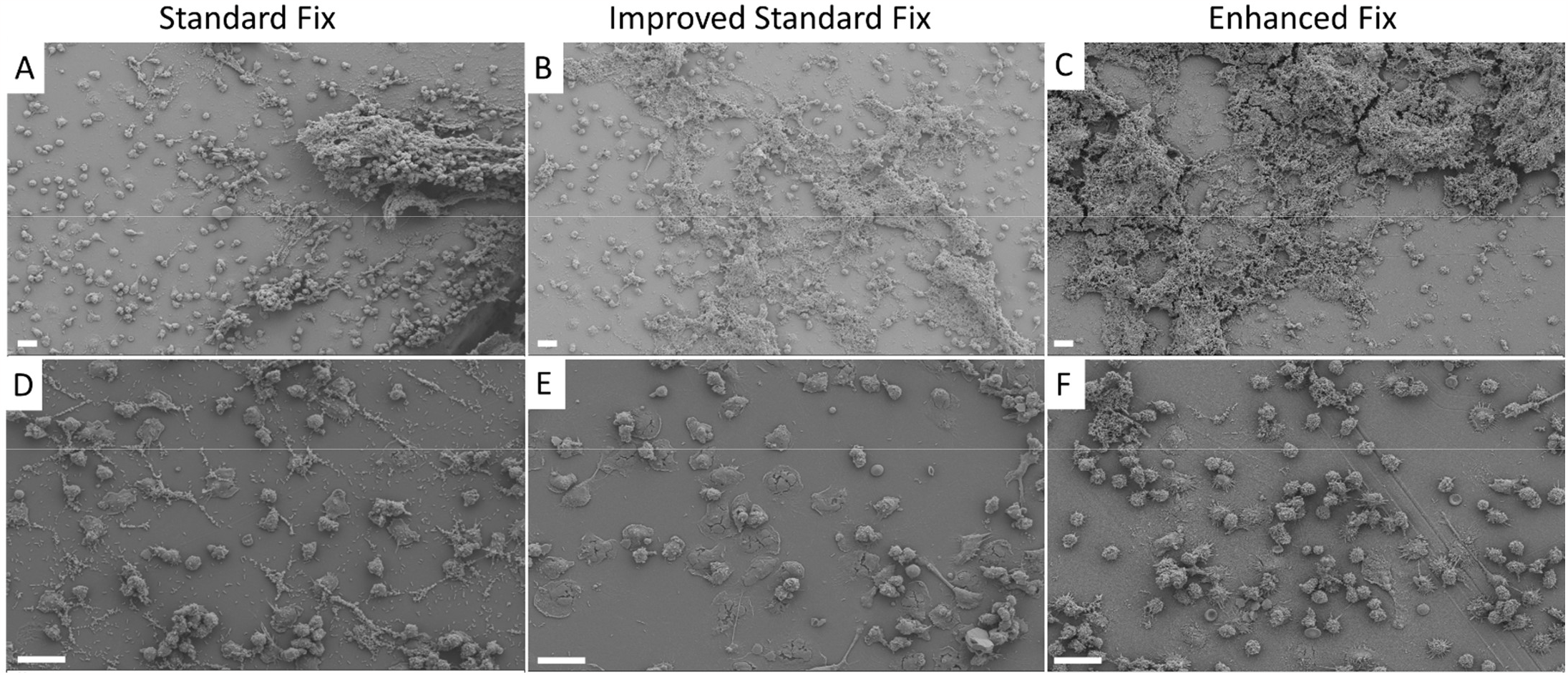
Representative SEM images of neutrophils interacting with Alginate+ biofilms for each of three fixation techniques. Panels D-F show higher magnification of the samples shown in A-C. Scale bars represent 20 µm.

Tannic acid has been previously reported to prevent shrinkage of neutrophils and allow them to retain their surface morphology [33, 40]. A good example of better surface morphology is given by neutrophils that appear to have employed the ‘frustrated’ phagocytosis mechanism. While spreading of the cell and characteristic fibril extensions are generally observed for all fixation methods, the enhanced fix best preserves these structures in three dimensions, as well as the wrinkled and ridged details of the neutrophils’ plasma membrane (Figure 3). We find that the retention of detail on the cell surface is substantially improved with the enhanced fix over that preserved by the other two fixes.

Furthermore, upon inspection of images from each fixation technique, the enhanced fix is the only one for which we observe the manifestation of a wide range of neutrophil responses. Figure 4 shows representative images of a neutrophil extracellular trap (A), degranulation (B), apoNETosis (C)[17], and neutrophil surface receptors at the interface with a biofilm (D). The fact that we do not observe these characteristics when the other two fixation methods are used indicates that standard fixation techniques may only preserve a fraction of neutrophil activity, and they are inadequate for detailed study of the complex immune response to infection.

**Figure 4.**
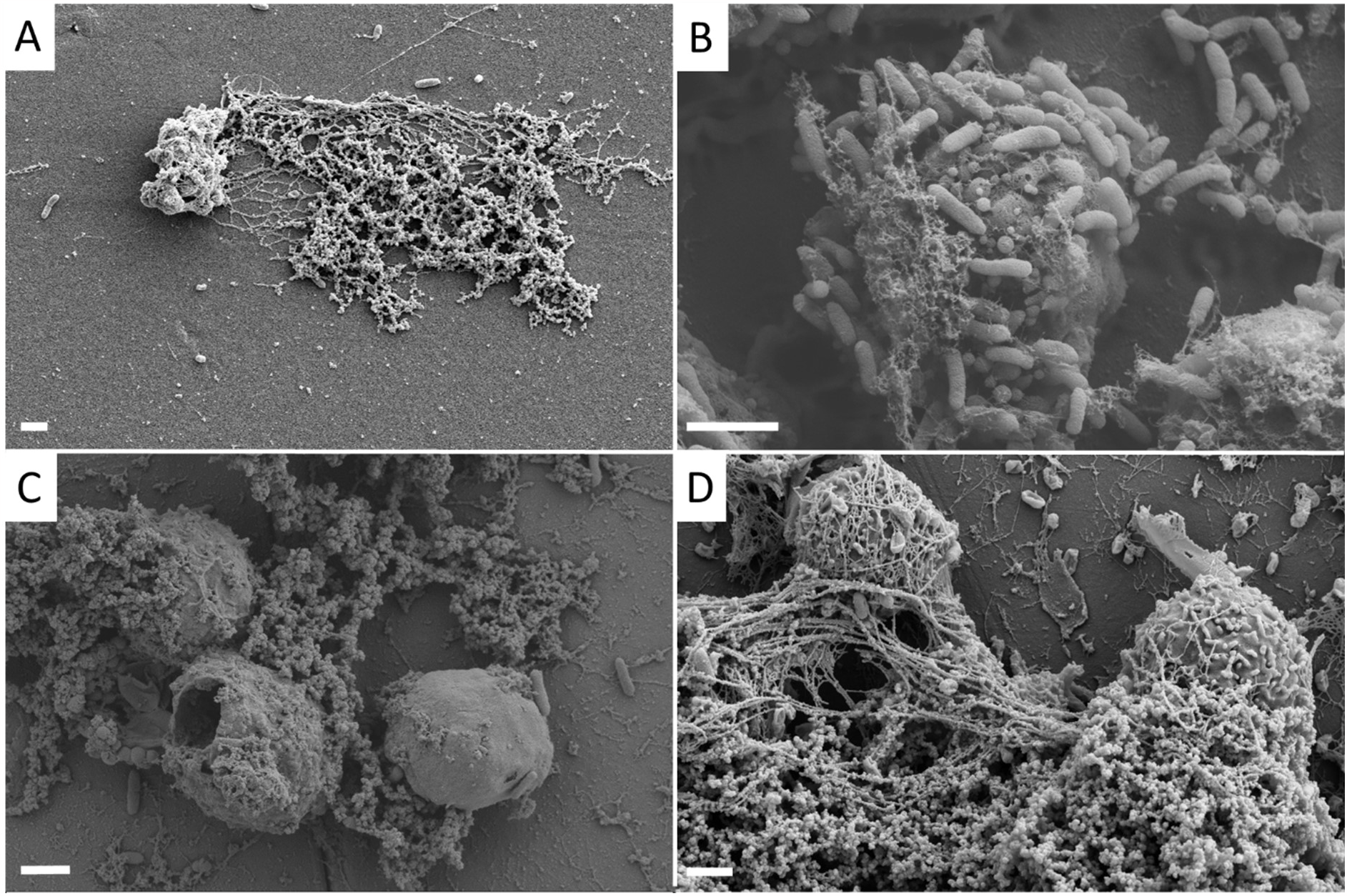
A) Optimized fixation preserves details of a neutrophil extracellular trap. B) A neutrophil releases antimicrobial granules to kill bacteria. C) Neutrophils may undergo apoptosis after releasing a NET, dubbed ‘apoNETosis’, leaving a ruptured cell membrane. D) Detailed surface topography of neutrophils interacting with a biofilm. Scale bars represent 2µm.

## Conclusion

In this study, we investigated three methods for fixing samples of biofilms and immune cells for SEM imaging of immune cell interactions with biofilms. We demonstrate that an enhanced fixative including the cationic dye alcian blue and tannic acid results in more successful preservation of EPS material, three-dimensional structure, and details of neutrophil killing mechanisms than do standard aldehyde-based fixatives. We observe higher yield of EPS material for both the WT and an alginate-overproducing strain of *P. aeruginosa*, notably when the matrix is dominantly composed of the anionic polysaccharide alginate. This preparation method also results in detailed visualization of neutrophil functional responses to biofilm infections, which may provide key insights for the development of biofilm-compromising treatments.

## Acknowledgements

SEM preparation and imaging was performed at the Center for Biomedical Research Support Microscopy and Imaging Facility at UT Austin (RRID:SCR_021756).

